# Microtubule bundling by MAP65-1 protects against severing by inhibiting the binding of katanin

**DOI:** 10.1101/520445

**Authors:** Graham M Burkart, Ram Dixit

## Abstract

The microtubule-severing enzyme katanin regulates the organization and turnover of microtubule arrays by the localized breakdown of microtubule polymers. In land plants, katanin (KTN1) activity is essential for the formation of linearly organized cortical microtubule arrays which determine the axis of cell expansion. Cell biological studies have shown that even though KTN1 binds to the sidewalls of single and bundled microtubules, severing activity is restricted to microtubule crossover and nucleation sites, indicating that cells contain protective mechanisms to prevent indiscriminate microtubule severing. Here, we show that the microtubule bundling protein MAP65-1 inhibits KTN1-mediated microtubule severing *in vitro*. Severing is inhibited at bundled microtubule segments and the severing rate of non-bundled microtubules is reduced by MAP65-1 in a concentration-dependent manner. Using various MAP65-1 mutant proteins, we demonstrate that efficient crosslinking of microtubules is crucial for this protective effect and that microtubule binding alone is not sufficient. Reduced severing due to microtubule bundling by MAP65-1 correlated to decreased binding of KTN1 to these microtubules. Taken together, our work reveals that crosslinking of microtubules by MAP65-1 confers resistance to severing by inhibiting the binding of katanin and identifies the structural features of MAP65-1 that are important for this activity.

**Highlight Summary:** Cortical microtubule bundles resist severing *in vivo*. Here, we show that crosslinking of microtubules by MAP65-1 inhibits severing in a dose-dependent manner by preventing katanin from binding to these microtubules.

## INTRODUCTION

The microtubule cytoskeleton is a multi-purpose structure whose configuration dynamically changes depending on the needs of the cell. Microtubule-severing enzymes have emerged as key players for the formation, remodeling and breakdown of microtubule arrays in eukaryotes (McNally and Roll-Mecak, 2018). Katanin was the first identified microtubule-severing enzyme and consists of a catalytic p60 subunit and a regulatory p80 subunit (McNally and Vale, 1993; Hartman et al., 1998). The p60 subunit belongs to the ATPases associated with diverse cellular activities superfamily and is sufficient for microtubule severing in an ATP-dependent manner (McNally and Vale, 1993; Hartman et al., 1998). The p80 subunit confers targeting and some degree of regulation of the p60 catalytic activity (Hartman et al., 1998; McNally et al., 2014; Wang et al., 2017). Biochemical and structural studies have shown that the p60 and p80 subunits assemble into hexameric ring structures which interact with the C-terminal tails of tubulin and remove tubulin dimers from microtubule sidewalls to promote severing (McNally and Vale, 1993; Hartman and Vale 1999; Zehr et al., 2017, Nithianantham, et al., 2018; Vemu, Szczesna, et al., 2018).

In the land plant *Arabidopsis thaliana*, the p60 subunit is encoded by a single gene called *KTN1* (Burk et al., 2001; Stoppin-Mellet et al., 2002) and null mutants exhibit developmental defects that are associated with aberrant organization of interphase and mitotic microtubule arrays (Bichet et al., 2001; Burk and Ye, 2002; Bouquin et al., 2003; Komis, Luptovčiak, Ovečka, et al., 2017). Live-imaging studies of the interphase cortical microtubule array revealed that KTN1 severs microtubules almost exclusively at microtubule crossover junctions and at nucleation sites to generate coaligned microtubule organization and to drive light-induced array reorientation (Wightman, Chomicki, et al., 2013; Nakamura et al., 2010; Zhang et al., 2013; Lindeboom et al., 2013; Fan et al., 2018). Interestingly, GFP-labeled KTN1 localizes not only to microtubule crossover and nucleation sites in a p80-dependent manner (Wang et al., 2017), but also localizes extensively along the shaft of cortical microtubules (Zhang et al., 2103; Lindeboom et al., 2013). The lack of severing along the cortical microtubule shaft suggests that off-target severing is inhibited, however the underlying mechanism remains unknown.

The neuronal microtubule-associated proteins tau and MAP2, which bundle and stabilize microtubules in axons and dendrites respectively, have been shown to protect microtubules against katanin activity when expressed in fibroblasts (Qiang, Yu, et al., 2006; Qiang et al., 2010). In addition, depletion of tau or phosphorylation-induced dissociation of tau from microtubules has been correlated with increased katanin-based microtubule severing in neurons (Qiang, Yu, et al., 2006). These data suggest that microtubule-crosslinking proteins inhibit microtubule severing by katanin.

Microtubule bundling is also a prominent feature of the plant cortical microtubule cytoskeleton and is thought to be important for the construction of higher-order microtubule structures (Shaw et al., 2003; Dixit and Cyr, 2004). Therefore, protecting cortical microtubule bundles against severing is likely to be important to maintain array integrity. Plant cortical microtubule bundling is mediated by the evolutionarily conserved MAP65 family of microtubule crosslinking proteins (Smertenko et al., 2004; Chang, Smertenko, et al., 2005; Smertenko, Chang, et al., 2006; Lucas et al., 2011; Lucas and Shaw 2012). Similar to the mammalian PRC1 and fungal Ase1p homologs, plant MAP65 crosslinks microtubules and dynamically labels the bundled regions of microtubules both *in vivo* (Lucas et al., 2011) and *in vitro* (Tulin et al., 2012), making it ideally suited to protect bundled regions of cortical microtubules against severing.

Here, we report that purified MAP65-1 protein inhibits microtubule severing *in vitro* in a concentration-dependent manner and that this effect is primarily due to the crosslinking of microtubules by MAP65-1 rather than just the binding of MAP65-1 to microtubules. In addition, we show that the inhibitory effect of MAP65-1 correlates with reduced binding of KTN1 to bundled microtubules. Together, our findings provide a mechanistic model for why bundled microtubules are not prone to severing in cells.

## RESULTS AND DISCUSSION

### GFP-KTN1 dwell time is reduced on bundled microtubules

KTN1 has been reported to bind to cortical microtubule bundles without detectably severing them (Zhang et al., 2013; Lindeboom et al., 2013). The likelihood of severing correlates to the dwell time of KTN1 on a microtubule, with the average dwell time for severing at crossover sites approximately 40 s (Zhang et al., 2013). We wondered whether microtubule bundles impair the binding of KTN1. To determine the dwell time of KTN1 on bundled microtubules, we conducted time-lapse imaging of *Arabidopsis thaliana ktn1-2* mutant plants co-expressing GFP-KTN1 (Lindeboom et al., 2013) and an mRuby-TUB6 microtubule marker. From these time series, we measured the duration of individual GFP-KTN1 puncta on single cortical microtubules, bundled cortical microtubules and at microtubule crossover sites (Figure 1, A and B). We found that the dwell time of GFP-KTN1 on bundled microtubules (4.2 ± 1.9 s) is dramatically lower than at crossover sites which are severed (49.7 ± 8.7 s) (Figure 1C). In addition, the dwell time of GFP-KTN1 on single microtubules which are not severed (4.0 ± 1.7 s) is comparable to that of bundled microtubules (Figure 1C).

**Figure 1.**
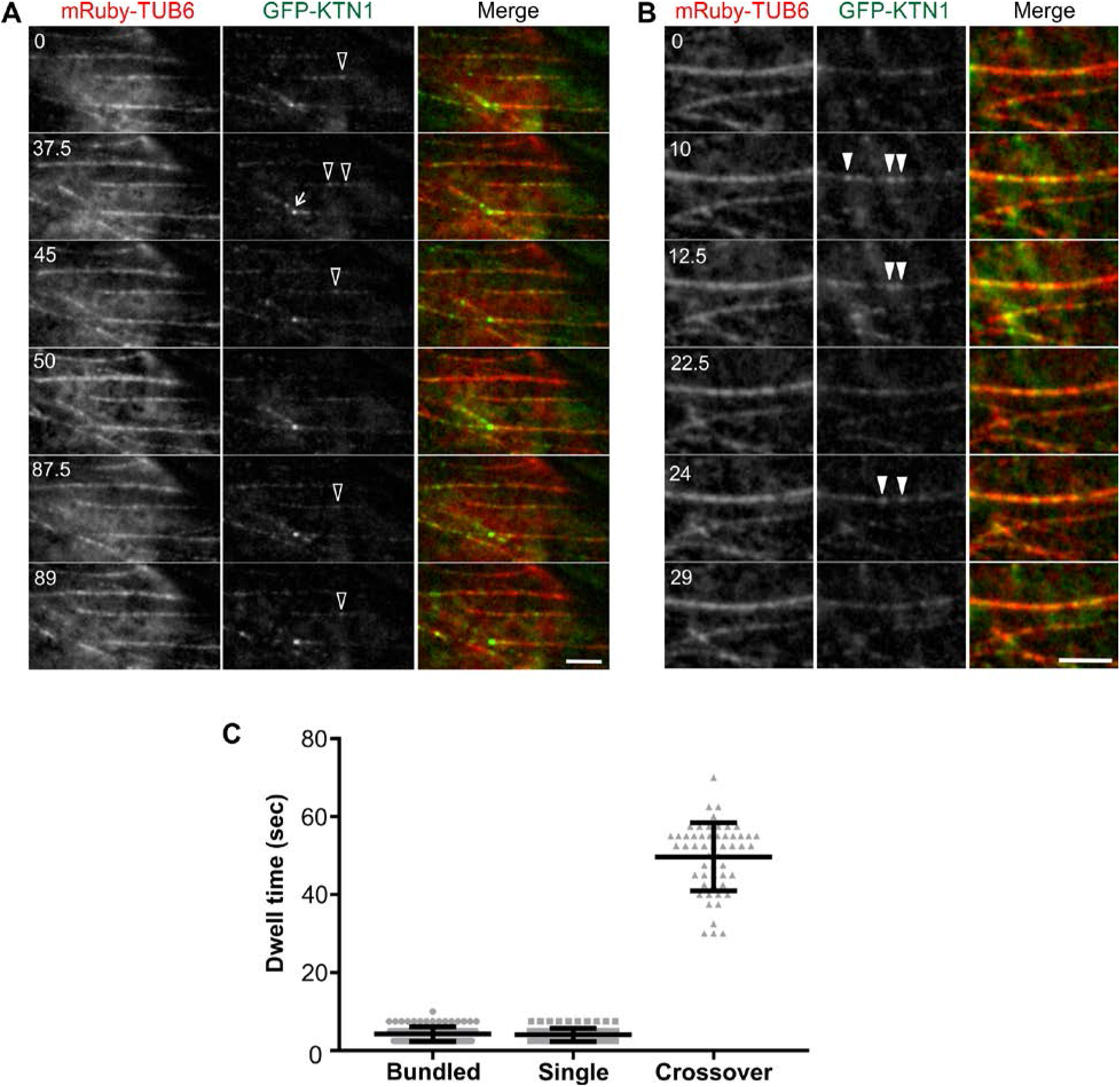
GFP-KTN1 binds more transiently to bundled cortical microtubules than to crossover sites. (A, B) Time course of GFP-KTN1 binding to single (A) and bundled (B) cortical microtubules *in vivo*. Empty arrowheads indicate GFP-KTN1 particles on single microtubules. Solid arrowheads indicate GFP-KTN1 particles on bundled microtubules. Arrow points to a crossover site. Scale bar is 5 μm. Numbers indicate time in seconds. (C) Dwell times of individual GFP-KTN1 particles on single microtubules (n = 96), bundled microtubules (n = 109), and at crossover sites (n = 54). Values are means ± s.d.

### The MAP65-1 protein inhibits KTN1-mediated microtubule severing

Proteins of the MAP65 family mediate bundling of cortical microtubules. In this study, we selected the MAP65-1 isoform because it abundantly binds to and bundles cortical microtubules *in vivo* (Smertenko et al., 2004; Gaillard, Neumann, et al., 2008; Lucas et al., 2011) and also bundles microtubules *in vitro* with similar characteristics (Tulin et al., 2012). To determine whether microtubule bundles created by MAP65-1 are resistant to KTN1 activity, we used an *in vitro* microtubule severing assay with purified recombinant KTN1 and MAP65-1 proteins (Figure 2A). Our purified KTN1 protein severs taxol-stabilized microtubules in an ATP-dependent manner (Figure 2, B and C, Movie 1) as reported previously (McNally and Vale, 1993; Stoppin-Mellet et al., 2002). We measured the microtubule fluorescence intensity over time in these assays and found that KTN1 severs essentially all microtubules by 6 minutes under our experimental conditions. Next, we pre-incubated microtubules with MAP65-1 for 10 minutes to induce bundling followed by the addition of KTN1 (Figure 2D, Movie 2). We found that microtubule severing was inhibited by MAP65-1 in a concentration dependent manner (Figure 2E). At 100nM MAP65-1, single microtubules and occasionally also bundled microtubules are severed (Figure 2D, solid arrowheads). However, at higher MAP65-1 concentrations both single and bundled microtubules are increasingly more intact, with severing essentially completed inhibited at 300nM MAP65-1 except for a few single microtubules (Figure 2D, empty arrowheads). These findings suggest that MAP65-1 can protect both single and bundled microtubules in a concentration-dependent manner, although the efficacy of protecting bundled microtubules is higher at any given concentration.

**Figure 2.**
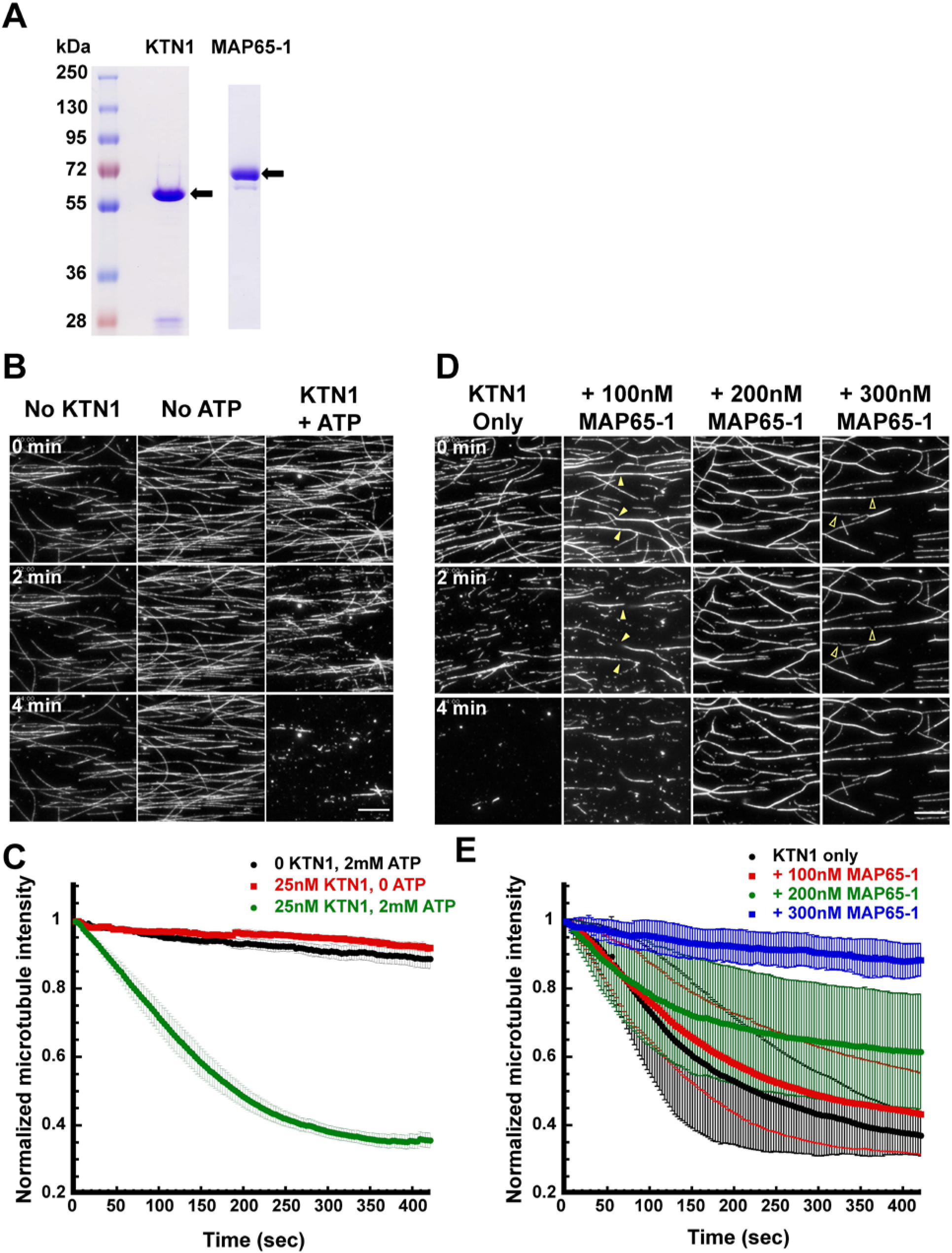
Microtubule bundling by MAP65-1 inhibits KTN1-mediated severing in a dose-dependent manner. (A) Coomassie Blue stained SDS-PAGE gel loaded with purified KTN1 or MAP65-1 proteins. Arrows indicate the expected bands for KTN1 and MAP65-1. (B) Time course of rhodamine-labeled, taxol-stabilized microtubules in the presence of either 2mM ATP, 25nM KTN1, or 25nM KTN1 and 2mM ATP. Scale bar is 10 μm. (C) Plot of microtubule signal intensity over time from experiments shown in (B). Each image in a series is normalized to the fluorescence intensity of the first frame of that series. Error bars represent s.e.m. of at least 4 separate protein preparations. (0 KTN1, 2mM ATP, n = 12 movies; 25nM KTN1, 0 ATP, n = 12 movies; 25nM KTN1, 2mM ATP, n = 18 movies). (D) Time course of rhodamine-labeled, taxol-stabilized microtubules in the presence of either 25nM KTN1 alone or with a 10-minute preincubation with 100nM, 200nM or 300nM of MAP65-1. Solid arrowheads indicate severing of bundled microtubules and empty arrowheads indicate severing of individual microtubules. (E) Plot of microtubule signal intensity over time from experiments shown in (D). Each image in a series is normalized to the fluorescence intensity of the first frame of that series. Error bars represent standard deviation. (25nM KTN1 alone, n = 8 movies; + 100nM MAP65-1, n = 10 movies; + 200nM MAP65-1, n = 16 movies; + 300nM MAP65-1, n = 10 movies).

### Microtubule crosslinking by MAP65-1 is required for efficient inhibition of severing activity

One possible mechanism by which MAP65-1 inhibits microtubule severing is by competing for the same binding site on the microtubule as KTN1. Alternatively, the cross-bridges formed by MAP65-1 between microtubules might sterically block KTN1 from accessing the microtubule. The first mechanism predicts that the microtubule-binding domain of MAP65-1 should be sufficient to inhibit microtubule severing. The second mechanism predicts that the crossbridge length of MAP65-1 would impact severing activity.

To test these predictions, we generated a novel MAP65-1 mutant protein that we refer to as MAP65-1 ND which consists only of the microtubule-binding domain and therefore cannot homodimerize to form microtubule bundles (Figure 3A). To study the contribution of the cross-bridge length, we took advantage of previously characterized MAP65-1 mutants (Tulin et al., 2012) that either decrease the spacing between microtubules (MAP65-1 ΔR1) or increase the spacing between microtubules (MAP65-1 R1R4) compared to wild-type MAP65-1 (WT MAP65-1), respectively (Figure 3A). To confirm that MAP65-1 ND does not bundle microtubules, we incubated WT MAP65-1, MAP65-1 ΔR1, MAP65-1 R1R4 and MAP65-1 ND with 500nM of rhodamine-labeled, taxol-stabilized microtubules for 10 minutes before imaging. We observed microtubule bundling with the wild-type, ΔR1, and R1R4 mutants but not with MAP65-1 ND mutant (Figure 3B). We also found that the MAP65-1 ΔR1 and MAP65-1 R1R4 mutants exhibited weaker and stronger microtubule bundling compared to WT MAP65-1, respectively (Figure 3B), probably because the longer rod domain of R1R4 bundles microtubules more efficiently compared to the shorter rod domain of ΔR1 (Tulin et al., 2012).

**Figure 3.**
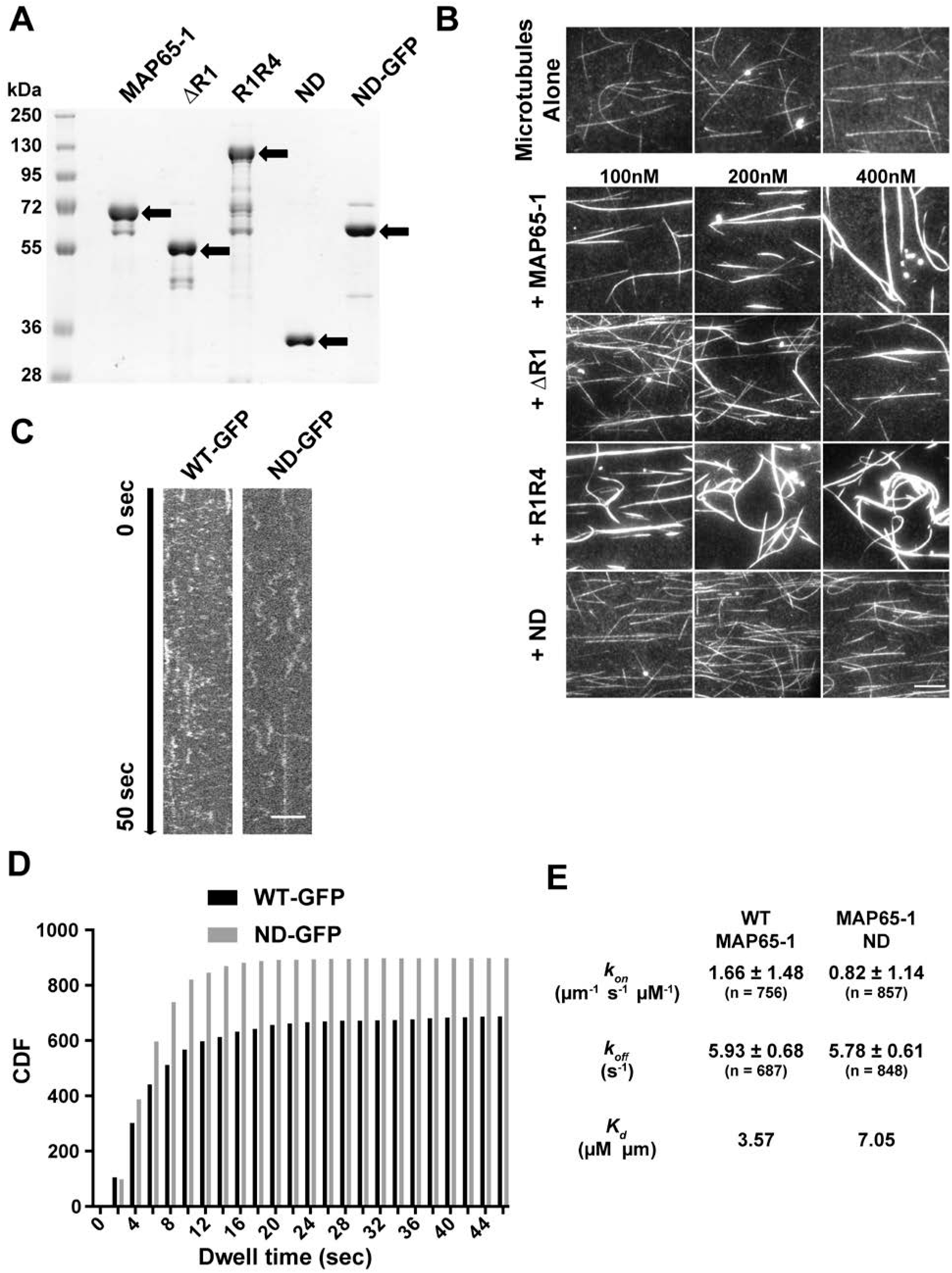
The MAP65-1 ND mutant does not bundle microtubules. (A) Coomassie Blue stained SDS-PAGE gel loaded with mutant versions of MAP65-1 protein. Arrows indicate the expected bands for each protein. (B) Fluorescence micrographs of rhodamine-labeled, taxol-stabilized microtubules alone or in the presence of varying concentrations of wild-type or mutant MAP65-1 proteins. All images were acquired under identical conditions. Scale bar is 10 μm. (C) Kymographs showing the behavior of WT MAP65-1-GFP and MAP65-1 ND-GFP on single microtubules. The brightness and contrast of the images were adjusted to better visualize the individual particles. Scale bar is 4 μm. (D) Cumulative distribution frequency plots of the dwell times of single WT MAP65-1-GFP (black) and MAP65-1 ND-GFP (gray) molecules. (E) Binding rate constant (*k_on_*), dissociation rate constant (k_*off*_), the apparent dissociation constant (K_d_) of WT MAP65-1-GFP and MAP65-1 ND-GFP determined from kymograph analysis of single molecules. Values represent mean ± s.d. (n = number of molecules).

Since we did not observe microtubule bundling with the MAP65-1 ND mutant, we wanted to confirm that this mutant protein binds to microtubules. We generated a MAP65-1 ND-GFP fusion protein and compared its microtubule binding to WT MAP65-1 GFP using single-molecule imaging conditions (Figure 3, C and D). We found that the binding rate constant (*k_on_*) of MAP65-1 ND was ~2-fold lower than that of WT MAP65-1 (Figure 3E) whereas the dissociation rate constant (*k_off_*) of MAP65-1 ND was similar to that of WT MAP65-1 (Figure 3E). Therefore, the apparent dissociation constant, *K_d_*, of MAP65-1 ND is about ~2-fold lower than that of WT MAP65-1. Based on these data we conclude that MAP65-1 ND binds to microtubules but with a slightly lower affinity compared to WT MAP65-1.

To examine the impact of the various MAP65-1 mutant proteins on katanin-mediated microtubule severing, we pre-incubated microtubules for 10 minutes with 200nM of WT or mutant MAP65-1 proteins before introducing KTN1. We found that the MAP65-1 ND mutant had a minimal inhibitory effect on microtubule severing compared to WT MAP65-1 (Figure 4, A and B), indicating that MAP65-1 probably does not directly compete with KTN1 for microtubule binding sites. In contrast, both MAP65-1 ΔR1 and MAP65-1 R1R4 reduced microtubule severing, underscoring that bundling is required to protect microtubules against severing. Furthermore, the degree of protection by the MAP65-1 mutants correlated with their strength of microtubule bundling (Figure 4, A and B).

**Figure 4.**
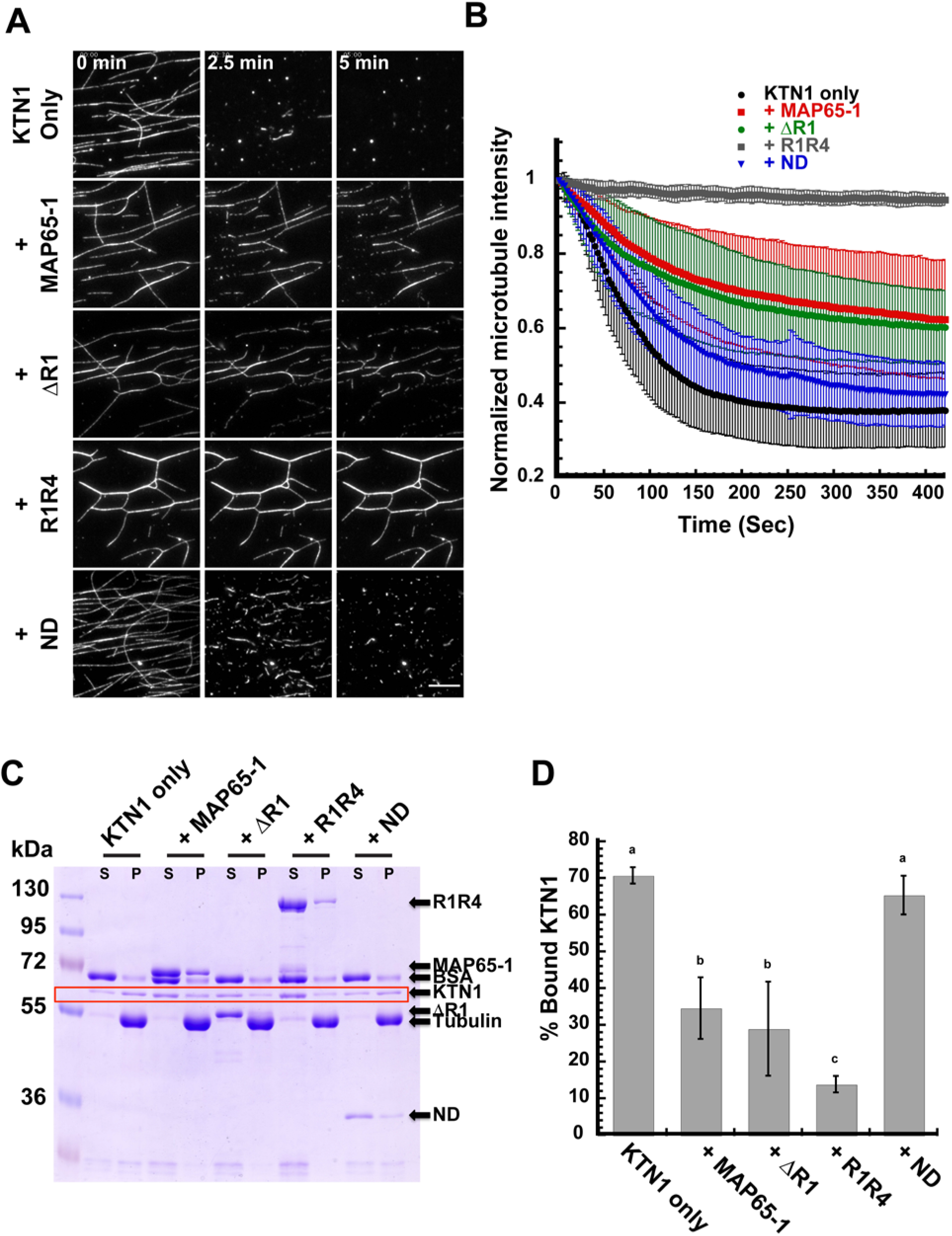
Bundling of microtubules by MAP65-1 is required to inhibit severing by decreasing the binding of KTN1 to microtubules. (A) Time course of rhodamine-labeled, taxol-stabilized microtubules in the presence of 25nM KTN1 alone or with a 10-minute pre-incubation with 200nM of either wild-type MAP65-1, MAP65-1 ΔR1, MAP65-1 R1R4, or MAP65-1 ND proteins. Scale bar is 10 μm. (B) Plot of microtubule signal intensity over time from experiments shown in (A). Each image in a series is normalized to the fluorescence intensity of the first frame of that series. Error bars represent standard deviation. (25nM KTN1, n = 10 movies; wild-type MAP65-1, n = 10 movies; MAP65-1 ΔR1, n = 12 movies; MAP65-1 R1R4, n = 10 movies; MAP65-1 ND, n = 10 movies). (C) Coomassie Blue stained SDS-PAGE gel of a microtubule co-sedimentation assay. 1μM rhodamine-labeled, taxol-stabilized microtubules were incubated with either 1μM KTN1 alone or were pre-incubated for 10 minutes with 2μM of the indicated MAP65-1 proteins before adding KTN1. S, supernatant; P, pellet. Arrows identify the different protein bands. The KTN1 bands are boxed in red. (D) Quantification of the microtubule-bound KTN1 fraction for the experiments shown in (C). The bars represent the mean ± s.d. (n = 6 for each condition). Letters above the bars indicate statistical groups with α < 0.05 using ANOVA.

### Microtubule bundling by MAP65-1 inhibits KTN1 binding

Based on our live imaging observation of reduced GFP-KTN1 dwell time on cortical microtubule bundles and our finding that microtubule bundling inhibits severing *in vitro*, we hypothesized that microtubule bundling by MAP65-1 inhibits the binding of KTN1 to microtubules. To test this, we performed microtubule co-sedimentation experiments. Microtubules were pre-incubated with either WT MAP65-1, MAP65-1 mutants, or with buffer alone for 10 minutes before introducing KTN1 (Figure 4C). We conducted these assays using the nonhydrolyzable ATP analog, AMPPNP, to stabilize the binding of KTN1 to microtubules without severing them. We found that the microtubule bound fraction of KTN1 decreased by 2-fold when microtubules were pre-incubated with either WT MAP65-1 or the MAP65-1 ΔR1 mutant (Figure 4D). In contrast, the MAP65-1 R1R4 mutant inhibited KTN1 binding by about 5-fold compared to the control. Notably, the MAP65-1 ND mutant did not significantly inhibit KTN1 binding to microtubules (Figure 4D). Based on these results we conclude that microtubule bundling inhibits severing by decreasing the binding of KTN1 to those microtubules.

Together, our findings indicate that bundling of microtubules by MAP65-1 is an effective protective mechanism to prevent unwanted microtubule severing by KTN1. Interestingly, the degree to which MAP65-1 counteracts KTN1 is significantly reduced when we used only the microtubule-binding domain of MAP65-1. One reason for this could be that the affinity of the microtubule-binding domain alone for microtubules is less compared to full-length MAP65-1. In addition, full-length MAP65-1 stays bound to microtubules for a significantly longer duration upon a bundling reaction (Tulin et al., 2012), whereas the microtubule-binding domain alone lacks this ability because it cannot engage in a bivalent interaction with two microtubules. In any case, our results reveal that the microtubule-binding domain alone is not sufficient to inhibit severing, consistent with our *in vitro* observation that single microtubules bound by MAP65-1 are more prone to severing than microtubules bundled by MAP65-1. Our examination of the impact of the rod domain on severing activity demonstrated a strong correlation between the strength of microtubule bundling and the degree of inhibition of microtubule severing. Therefore, the ability to crosslink microtubules is a major determinant of the protective effect of MAP65-1.

Recent work has discovered that at short timescales katanin damages microtubules by removing tubulin dimers from microtubule sidewalls (Vemu, Szczesna, et al., 2018). These injury sites can be healed by the incorporation of fresh GTP-tubulin dimers. However, prolonged incubation with katanin tips the balance towards damage and leads to microtubule severing (Vemu, Szczesna, et al., 2018). Since the residence time of KTN1 is lower on bundled microtubules, one potential mechanism for why bundled microtubules are more resistant to severing is a reduction in the rate of removal of tubulin dimers from these microtubules. If the dimer removal rate is slower than the rate of insertion of new GTP-tubulin dimers, then the microtubule lattice is more likely to be repaired than get severed. Another potential mechanism that is not mutually exclusive is that MAP65-1 crosslinks brace damaged microtubules and thus inhibit complete severing from occurring in these regions.

Our finding that MAP65-1 inhibits severing in a concentration-dependent manner suggests a potential mechanism for cells to tune the level of protection against KTN1 activity. For example, cells can regulate the amount of MAP65-1 protein to globally modulate the extent of microtubule severing. Furthermore, phosphorylation of MAP65-1, which is known to decrease its microtubule binding (Smertenko, Chang, et al., 2006), could serve as a mechanism to locally control the extent of microtubule severing. Therefore, the balance between bundling and severing provides a potential mechanism to control the architecture and stability of microtubule arrays.

The *MAP65* gene family has expanded considerably in land plants compared to mammals and yeast (Hussey et al., 2002). Plant MAP65 isoforms differ considerably in polypeptide length and in their preference for polarity of microtubule bundling (Gaillard, Neumann, et al., 2008; Smertenko et al., 2008; Fache et al., 2010; Ho et al., 2012). Certain plant MAP65 isoforms also localize to specific microtubule arrays during the cell cycle or even to specific zones of particular microtubule arrays (Van Damme et al., 2004; Li, Sun, et al., 2017). Future work will determine whether different MAP65 isoforms differ in their ability to inhibit severing, which would provide another mechanism for plants to differentially protect microtubule arrays from severing. Additional work is also needed to determine whether the mammalian and yeast homologs of MAP65-1 inhibit microtubule severing in the mitotic microtubule arrays to which they localize.

## MATERIALS AND METHODS

### Plant materials, growth conditions and imaging

We generated a pGAG UBQ10promoter::mRuby-TUB6-Kan^R^ plasmid using multisite-Gateway recombination and introduced it into *Arabidopsis thaliana* Col-0 plants via *Agrobacterium tumefaciens*-mediated transformation. Homozygous transformants were crossed to the *ktn1-2* mutant expressing GFP-KTN1. Seeds from plants homozygous for the *ktn1-2* mutation and at least heterozygous for GFP-KTN1 and mRuby-TUB6 were surface sterilized, plated on 1/2X Murashige and Skoog with 1% agar and stratified in the dark for 2 days at 4°C. Seedlings were grown at 22°C in a 16-hour/8-hour light/dark cycle for 3-4 days before observation. Seedlings were imaged by VAEM microscopy (Konopka and Bednarek, 2008) using 488 nm and 514 nm laser lines for excitation and 500-550 nm and 582-636 emission filter sets for GFP and mRuby imaging, respectively. Two-color, time-lapse images were collected every 2.5 seconds. Image analysis was conducted using the Fiji ImageJ package (Schindelin et al., 2012).

### Constructs for recombinant protein expression

pMAL C5-X (New England Biolabs) was modified by using PCR to add a 6x-histidine tag in frame upstream of the maltose binding protein (MBP) coding sequence and a tobacco etch virus protease (TEV) cleavage site in frame downstream of the MBP coding sequence, generating the expression vector pHis-MAL-TEV (pHMT). The cDNA sequence of KTN1 was introduced into pHMT using the NotI and SalI restriction sites.

The C-terminal microtubule-binding domain of MAP65-1 (amino acids 369-616) was cloned by PCR from the pTEV-MAP65-1 construct (Tulin et al., 2012) with primers CATATGTCTGCTCGTGAGAGAATCATGT and GGATCCTCATGGTGAAGCTGGAAC introducing NdeI and BamHI restriction sites for cloning into pTEV. To generate the ND-GFP construct, primers CATATGTCTGCTCGTGAGAGAATCATGT and GGATCCTGGTGAAGCTGGAACTTG were used to amplify the MAP65-1 ND fragment, and primers GGATCCATGGTGAGCAAGGGCGAG and GAGCTCTTACTTGTACAGCTCGTC were used to amplify mEGFP. The PCR products were both digested with BamHI and ligated together. The ligation was then used as a template for PCR using primers CATATGTCTGCTCGTGAGAGAATCATGT and GAGCTCTTACTTGTACAGCTCGTC. This PCR product was cloned into pTEV using NdeI and SacI. All constructs were confirmed by DNA sequencing.

### Expression and purification of recombinant KTN1 and MAP65-1 proteins

The pHMT-KTN1 plasmid was transformed into BL21-CodonPlus (DE3)-RIPL competent cells (Agilent Technologies). Protein was affinity-purified using Ni-NTA agarose (Qiagen #301210), and exchanged into severing buffer I (SBI) (pH 7.0, 20mM HEPES, 50mM NaCl, 75mM MgSO_4_, 2mM MgCl_2_, 10% sucrose, 5mM DTT, 50μM ATP) using PD-10 desalting columns (GE Healthcare # 17085101). Protein was then digested with TEV protease and run through a subtractive Ni-NTA column to remove TEV protease and uncut protein. MAP65-1 proteins were purified from Rosetta DE3 cells as described previously (Tulin et al., 2012). Protein aliquots were snap frozen in liquid nitrogen and stored at −80°C until use.

### Microtubule severing assays

Imaging chambers were prepared by affixing silanized coverslips to slides using strips of double-sided tape. A solution of 0.4% Anti-tubulin antibody (Sigma #T4026) in severing buffer II (SBII) (pH 7.0, 20mM HEPES, 3mM MgCl_2_, 10% sucrose) was coated on the coverslip and then blocked with a 5% pluronic F-127 (Sigma #P2443) solution in SBII. A 1:100 dilution of 5 mg/mL taxol-stabilized 1:25 rhodamine-labeled porcine microtubules was then flowed in and allowed to bind to the antibody for 5 minutes. Excess microtubules were then washed out with two washes of 20μM taxol in SBII. Severing mix (25nM KTN1, 2mM ATP, 50mM DTT, 800 μg/mL glucose oxidase, 175 μg/mL catalase, 22.5 mg/mL glucose, 20μM taxol in SBII) was then flowed in and the imaging chamber was immediately mounted and imaged by total internal reflection fluorescence microscopy. Time-lapse images were acquired every 3 seconds for 7.5 minutes using a back-illuminated electron-multiplying CCD camera (ImageEM, Hamamatsu, Bridgewater, NJ) and a 582-636nm emission filter set. Image analysis was performed using the Fiji ImageJ package (Schindelin et al., 2012).

For severing assays conducted with MAP65-1 proteins, SBII with 20μM taxol and the indicated amount of MAP65-1 protein was flowed in following the microtubule washout step and incubated for ten minutes before the severing mix (supplemented with the same concentration of MAP65-1 protein used for the incubation step) was flowed in and then chamber was mounted and imaged as described above.

### Single molecule imaging

Imaging chambers were prepared as for the microtubule severing assays. Following the microtubule washout step, 2.5nM MAP65-1 or 10nM MAP65-1 ND-GFP were then flowed in and immediately imaged. An initial snapshot of the microtubules was taken using the 582-636nm emission filter set. Streaming-mode images were acquired with the 500-550 nm filter set every 100 milliseconds for 50 seconds. Kymographs were generated along the microtubules and analyzed using the Fiji ImageJ package (Schindelin et al., 2012). Bright, static particles were considered to be protein aggregates and were not included in our analysis.

### Microtubule bundling assays

500nM taxol-stabilized, 1:25 rhodamine-labeled microtubules were incubated for 10 minutes with the indicated concentrations of MAP65-1 proteins, then flowed into an imaging chamber and imaged using the 582-636 nm emission filter set. All images were acquired with identical settings so that signal intensity reflects the degree of microtubule bundling.

### Microtubule co-sedimentation assays

To assess KTN1 binding to microtubules in the presence of MAP65-1 mutant proteins, 1μM of rhodamine-labeled, taxol-stabilized microtubules were co-incubated with 2μM of MAP65-1 proteins for 10 minutes in SBII supplemented with 50μM DTT, 40μM taxol, 0.1 mg/mL BSA and 2mM AMPPNP before adding 1μM KTN1 to the mixture. This mixture was then incubated at room temperature for 30 minutes before centrifugation at 15°C for 25 minutes. The supernatant and pellet fractions were run out on SDS-PAGE and Coomassie stained. Band densitometry was performed using Fiji software (Schindelin et al., 2012). Statistical analysis was performed using KaleidaGraph (Synergy Software).

## ACKNOWLEDGEMENTS

We thank Dr. Magdalena Bezanilla (Dartmouth College) for providing the cDNA for mRuby and pDonor plasmids and Dr. David Ehrhardt (Carnegie Institute for Science, Stanford University) for providing the GFP-KTN1 complemented *ktn1-2* mutant. This work was supported by the National Institute of Health Grant R01 GM114678.

## Abbreviations

Ase1p,: anaphase spindle elongation protein 1;
GFP,: green fluorescent protein;
KTN1,: katanin;
MAP2,: microtubule-associated protein 2;
MAP65,: microtubule-associated protein of 65 kDa size;
PRC1,: protein regulator of cytokinesis 1.

